# Membrane-dependent structural organization of cowpox virus CPXV012 and its recognition of TAP

**DOI:** 10.64898/2026.06.12.731803

**Authors:** Natalia Karska, Behnaz Mizraeli, Magdalena J. Ślusarz, Przemysław Karpowicz, Igor Zhukov, Sylwia Rodziewicz-Motowidło

## Abstract

Cowpox virus CPXV012 inhibits MHC class I antigen presentation by interfering with TAP-dependent peptide transport, but its membrane-dependent structural organization and dynamic behavior remain incompletely defined. Here, we investigated the conformational properties of CPXV012 in membrane-mimicking environments and in a model of the CPXV012–TAP complex. CPXV012 was divided into three peptide constructs corresponding to the N-terminal cytosolic region, transmembrane segment, and C-terminal ER-luminal domain. The peptides were analyzed by circular dichroism spectroscopy, multidimensional NMR spectroscopy, and molecular dynamics simulations, and the resulting structural information was integrated into a full-length CPXV012 model.

CD spectra showed that CPX-E1 and CPX-C2 are predominantly disordered in aqueous solution but acquire ordered, mainly α-helical features in DPC micelles. NMR analysis in DPC-d38 micelles provided residue-level assignments and structural restraints supporting restrained structure calculations for both peptides. In three independent 1 µs molecular dynamics simulations of the CPXV012–TAP complex, CPXV012 preserved a reproducible two-helical organization. The N-terminal/transmembrane region behaved as a relatively stable structural element, whereas the ER-luminal segment showed greater local flexibility. Interface analysis indicated that CPXV012 contacts both TAP1 and TAP2, with recurrent interactions concentrated in the luminal Y47–I69 region and involving polar and charge-complementary contacts.

These results support a model in which membrane-associated structuring positions CPXV012 for TAP recognition, while the flexible ER-luminal region forms the main TAP-interacting surface. This structural framework complements existing functional models of CPXV012-mediated TAP inhibition.

## Introduction

Cowpox virus interferes with MHC I antigen presentation through at least two mechanistically distinct proteins. CPXV203 retains fully assembled MHC I molecules in the ER/Golgi pathway by exploiting a KDEL-receptor-dependent retrieval mechanism [Byun et al., 2007; McCoy et al., 2012], **whereas** CPXV012, also referred to as CPXV12 in earlier studies, inhibits TAP-dependent peptide translocation into the ER [Alzhanova et al., 2009; Luteijn et al., 2014]. Deletion studies showed that CPXV012 and CPXV203 act independently but cooperatively, and that removal of both proteins restores MHC I surface expression and T-cell stimulation in CPXV-infected cells [Alzhanova et al., 2009]. CPXV012 was subsequently characterized as a small type II ER membrane protein that **blocks** TAP-mediated peptide transport by preventing ATP binding to TAP [Luteijn et al., 2014]. Mechanistic studies further showed that CPXV012 does not primarily inhibit peptide binding to TAP, but acts after peptide binding and peptide-induced conformational rearrangement, consistent with a lumenal trans-inhibition mechanism mediated by its C-terminal ER-luminal region [Lin et al., 2014].

More recently, cryo-EM structures of TAP in complex with viral inhibitors, including CPXV012, provided structural snapshots of inhibited TAP states and showed that structurally diverse viral proteins can converge on a shared strategy of stalling the TAP transport cycle [Lee et al., 2025]. However, these structures describe the TAP-bound state and do not fully define **the** intrinsic conformational behavior of isolated CPXV012 in a membrane-like environment. In particular, the structural properties of the N-terminal cytosolic segment, the transmembrane region, and the ER-luminal domain remain incompletely understood outside the TAP complex. This gap is relevant because CPXV012 is a small hydrophobic membrane protein whose inhibitory region is spatially constrained by membrane anchoring and may depend on the lipid environment for its organization and TAP recognition [Luteijn et al., 2014; Lin et al., 2014].

Although the TAP-bound state of CPXV012 has recently been structurally characterized, the intrinsic membrane-dependent conformational properties of CPXV012 and its individual regions remain insufficiently understood. In particular, it remains unclear how the cytosolic, transmembrane, and ER-luminal regions of CPXV012 are locally organized in membrane-mimicking environments and how this local organization relates to the dynamic behavior of full-length CPXV012 in the context of open-conformation TAP.

In this study, we investigated the structural and conformational properties of CPXV012 in membrane-mimicking environments using circular dichroism spectroscopy, multidimensional NMR spectroscopy, and molecular dynamics simulations. To obtain domain-specific structural information, CPXV012 was divided into three peptide constructs corresponding to its N-terminal cytosolic segment, transmembrane region, and C-terminal ER-luminal domain. The peptide-derived structural information was then used to support modeling of full-length CPXV012 in an endoplasmic reticulum-mimetic membrane environment and to explore its possible orientation relative to the TAP complex. This approach provides a membrane-oriented structural framework for CPXV012 and complements existing functional and structural studies of its TAP-bound inhibitory state.

## Results

### Design and synthesis of CPXV012-derived peptides

CPXV012 is a 69-residue type II ER membrane protein composed of three topological regions: a short N-terminal cytosolic segment, a central transmembrane segment, and a C-terminal ER-luminal domain involved in TAP inhibition [Luteijn et al., 2014; Lin et al., 2014]. Based on this topology, the CPXV012 sequence was divided into three peptide constructs corresponding to these regions: CPX-C2, comprising residues 1–12; CPX-T2, comprising residues 13–35; and CPX-E1, comprising residues 36–69 (Figure 1A). This design allowed the cytosolic, transmembrane, and ER-luminal parts of CPXV012 to be analyzed separately by CD spectroscopy, NMR spectroscopy, and molecular dynamics simulations.

**Figure 1.**
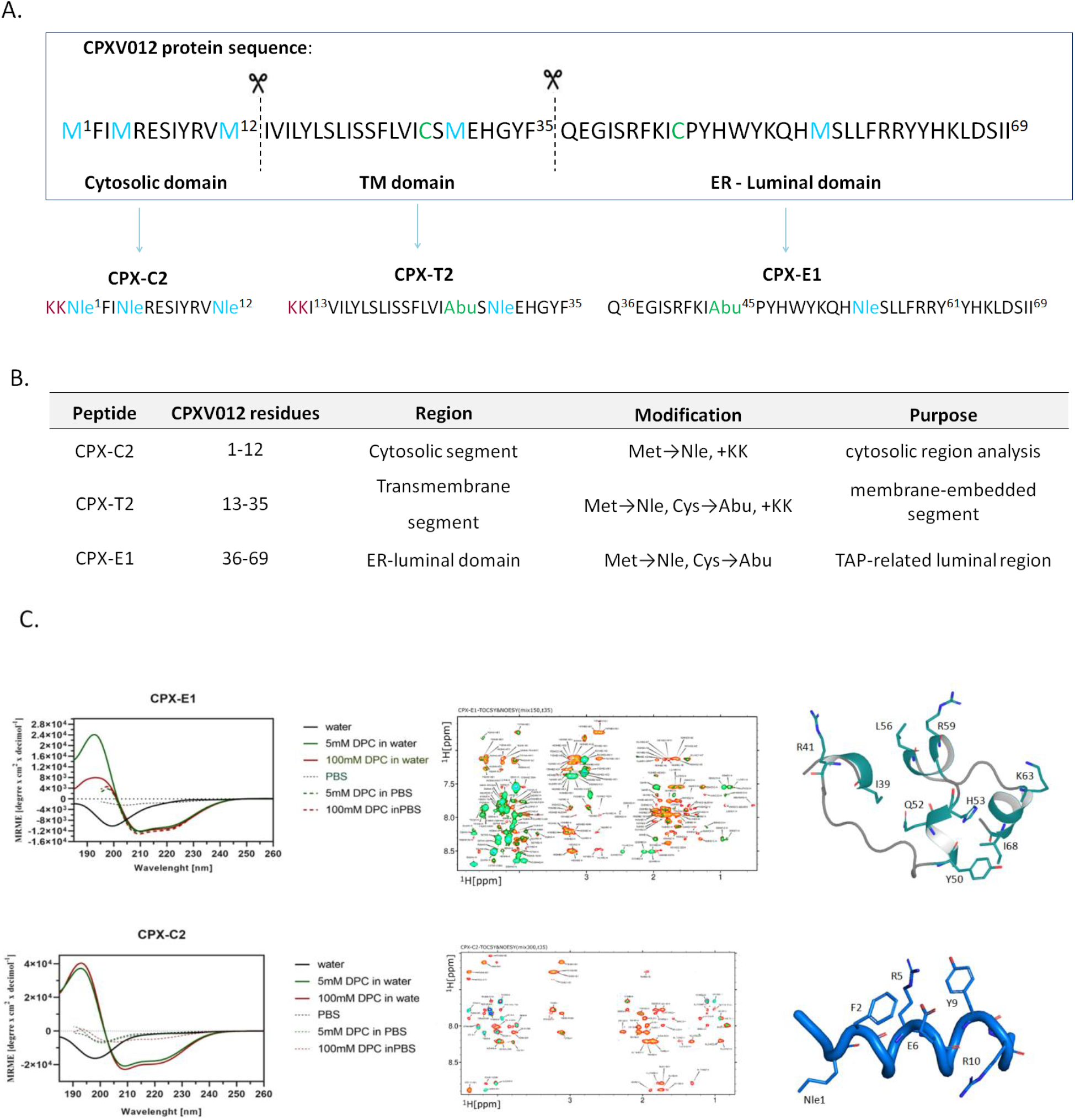
(A) Design and structural characterization of CPXV012-derived peptide constructs. **(A)** Schematic representation of the full-length CPXV012 sequence and the peptide segmentation strategy used for structural and biophysical analysis. CPXV012 was divided into three constructs corresponding to its predicted topological regions: CPX-C2, comprising the N-terminal cytosolic segment (residues 1–12); CPX-T2, corresponding to the transmembrane segment (residues 13–35); and CPX-E1, corresponding to the ER-luminal region involved in TAP recognition and inhibition (residues 36–69). Methionine and cysteine residues selected for substitution are indicated in the sequence. **(B)** Summary of the CPXV012-derived peptides used in this study, including residue range, corresponding protein region, introduced modifications, and experimental purpose. Methionine residues were replaced with norleucine (Nle), and cysteine residues were replaced with (S)-2-aminobutyric acid (Abu). The N-terminal Lys-Lys extension (KK), where present, was introduced to improve peptide solubility and is not part of the native CPXV012 sequence. **(C)** Biophysical and structural characterization of CPX-E1 and CPX-C2. The upper row shows data for the ER-luminal peptide CPX-E1, and the lower row shows data for the N-terminal cytosolic peptide CPX-C2. Left panels show far-UV circular dichroism spectra recorded in water, PBS, and DPC-containing conditions. Middle panels show overlaid TOCSY and NOESY spectra used for resonance assignment and identification of NMR-derived distance restraints. Right panels show NMR-derived structural models of CPX-E1 and CPX-C2 obtained from restrained structure calculations.

A peptide-based strategy was selected because full-length CPXV012 is a small hydrophobic membrane protein whose isolation in a homogeneous, tag-free form is challenging for high-resolution biophysical analysis. Similar fragment-based approaches have been successfully applied to structural studies of small viral membrane proteins [Karska et al., 2019]. The modular approach was used to determine local conformational propensities of individual CPXV012 regions under membrane-mimicking conditions. Because isolated peptides may not fully reproduce all long-range interactions present in the full-length protein, the fragment-derived structural information was subsequently integrated into a full-length CPXV012 model and evaluated in an endoplasmic reticulum-mimetic membrane environment by molecular dynamics simulations.

To improve chemical stability during synthesis and purification, methionine and cysteine residues were replaced with conservative isosteric analogues: norleucine and (S)-2-aminobutyric acid, respectively [Moroder, 2005]. The resulting CPXV012-derived peptides were obtained by solid-phase peptide synthesis, purified by RP-HPLC, and verified by mass spectrometry.

To facilitate structural investigations, the CPXV012 protein sequence was divided into three peptides: CPX-C2 comprising the N-terminal cytosolic region (residues 1–12), CPX-T2 representing the transmembrane segment (residues 13–35), and CPX-E1 corresponding to the ER-luminal domain (residues 36–69) (Figure 1). This modular design enabled separate analysis of membrane-associated and luminal structural determinants using CD spectroscopy, multidimensional NMR spectroscopy, and molecular dynamics simulations.

### Structural characterization of CPXV012-derived peptides in membrane-mimicking environments

Far-UV CD spectroscopy was used to evaluate the secondary-structure preferences of CPXV012-derived peptides in aqueous and membrane-mimicking environments. In water, CPX-E1 and CPX-C2 displayed spectra dominated by a negative band near 198–200 nm, consistent with predominantly disordered conformations. PBS alone produced only weak CD signals for both peptides.

Addition of DPC induced pronounced spectral changes. For **CPX-E1**, DPC in water produced a positive band near 192–195 nm and a broad negative region between approximately 208 and 225 nm, indicating increased helix-like structural organization. A similar negative region was also observed in DPC/PBS, although the positive band near 195 nm was less intense than in DPC/water. Thus, CPX-E1 acquired ordered conformational features in the presence of DPC, but the spectral response depended on both DPC concentration and buffer composition. CPX-C2 showed an even more pronounced response to DPC in water. In both 5 mM and 100 mM DPC, the spectra displayed a positive band near 193–195 nm and two negative bands around 208 and 222–225 nm, characteristic of increased α-helical content. The helical features were slightly stronger in 100 mM DPC than in 5 mM DPC. In contrast, the DPC-induced changes were substantially weaker in PBS, indicating that ionic strength or buffer composition reduced the interaction of CPX-C2 with the micellar environment.

Together, these results indicate that CPX-E1 and CPX-C2 are predominantly disordered in aqueous solution but acquire ordered, mainly helix-like conformational features in the presence of DPC micelles. The extent of this structural organization is peptide-specific and depends on the solvent conditions. These data show that DPC micelles promote ordered, predominantly helix-like conformational features in both CPX-E1 and CPX-C2, but the magnitude and concentration dependence of this effect differ between the two peptides and are modulated by buffer composition.

To obtain residue-level structural information, CPX-E1 and CPX-C2 were analyzed by NMR spectroscopy in DPC-d_38_ micelles. Complete ^1^H, ^15^N, and ^13^C resonance assignments were obtained for all residues of both peptides using homonuclear two-dimensional TOCSY and NOESY spectra complemented by heteronuclear ^1^H–^13^C and ^1^H–^15^N HSQC experiments. The NOESY spectra contained both sequential and medium-range contacts, supporting restrained structure calculations for both peptides. The 3D structure of the CPX-E1 was established on base 178 unique distance constraints, together with 53 restraints to ϕ, ψ, and χ1 torsion angles, and 10 constraints for hydrogen bonds (Table S1). The established high-resolution 3D structure presented as three helix motif comprised 310-helix (Y50–I54), α-helix2 (F58–Y61), and 310-helix3 (K64 –I68) (Figure 1C). Further inspection of evaluated structure reveals existence of specific motif formed by two histidines (H53, and H63), together with Y61. The calculated distances between majority of protons in side-chains of both histidines is less than 5.0 Å. The existence contacts between resides plays an important role to stabilize three helix motif in CPX-E1 peptide.

### Conformational behavior of CPXV012 in the open-conformation TAP model

To evaluate the conformational behavior of CPXV012 in the context of open-conformation TAP, three independent all-atom MD trajectories were performed for the CPXV012–TAP model. The final CPXV012 conformations extracted from the three trajectories showed preservation of the overall helical organization of the modeled protein, although the precise boundaries of the helical regions differed between simulations (Figure 2). In trajectory 1, helices were observed within residues M4–Q36 and P46–Y62. In trajectory 2, the main helical regions extended from M4 to E37 and from P46 to H63, with a short additional ordered segment at D66–I68. In trajectory 3, helices were observed within residues M1–F35 and P46–Y62. Overall, the N-terminal/transmembrane helical region was conserved across all trajectories, whereas the ER-luminal TAP-related region displayed greater variability, particularly toward its C-terminal end. These results indicate that CPXV012 maintains a reproducible global fold in the open-conformation TAP model while retaining local flexibility in the luminal segment implicated in TAP inhibition.

**Figure 2.**
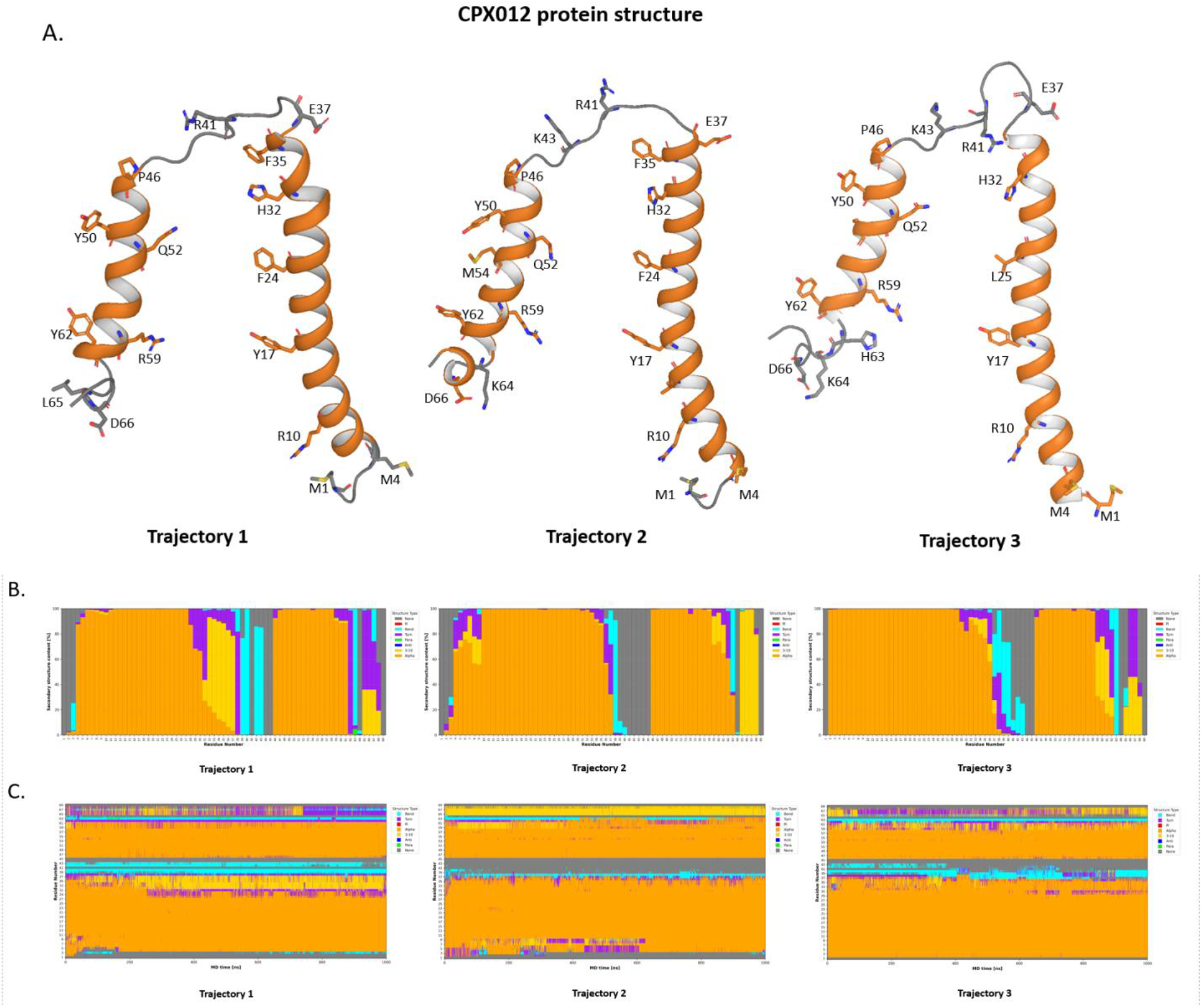
Final CPXV012 conformations obtained from three independent MD trajectories in the context of open-conformation TAP. A. Final conformations of CPXV012 obtained from three independent molecular dynamics trajectories using the open conformation of the TAP transporter as the structural reference. The CPXV012 backbone is shown in cartoon representation, and selected residues are displayed as sticks. The comparison of the three trajectories illustrates the reproducibility of the modeled CPXV012 fold and the trajectory-dependent flexibility of the TAP-related luminal segment. **The secondary structure analysis (DSSP) of CPXV012 in three trajectory with TAP during 1 µs molecular dynamics simulations: B**. Time-resolved DSSP secondary structure map for each residue over the simulation trajectory. **C**. Per-residue secondary structure composition expressed as percentage occupancy based on DSSP analysis.

DSSP analysis indicated that the N-terminal/transmembrane region of CPXV012 was the most reproducible structural element across the three trajectories. In trajectory 1, the main helical segments were assigned to residues M4–Q36 and P46–Y62. In trajectory 2, helicity extended from M4 to E37 and from P46 to H63, with an additional short ordered segment at D66–I68. In trajectory 3, the N-terminal/transmembrane helix extended from M1 to F35, whereas the luminal helix was observed between P46 and Y62. Thus, the N-terminal/transmembrane helical region was consistently preserved, while the ER-luminal region showed greater local variability, particularly at its C-terminal end.

The RMSF profiles further supported this two-region behavior. The lowest mobility was associated with the helical parts of CPXV012, whereas increased fluctuations were observed at the N-terminal residues and in the linker region around E37–K43. This linker separates the N-terminal/transmembrane helix from the ER-luminal TAP-interacting region and therefore appears to provide local conformational flexibility without disrupting the global organization of CPXV012. Time-resolved snapshots extracted every 100 ns confirmed that CPXV012 did not undergo unfolding during the simulations, but instead sampled related conformations in which the luminal segment remained mobile relative to the more stable N-terminal/transmembrane helix.

Together, these results indicate that CPXV012 adopts a reproducible two-helical organization in the open-conformation TAP model. The N-terminal/transmembrane region behaves as a relatively stable structural anchor, whereas the ER-luminal region retains local flexibility that may allow CPXV012 to maintain or adjust contacts with TAP during the simulation.

### CPXV012–TAP interface analysis

The representative final models of the CPXV012–TAP complex showed that CPXV012 associates with the TAP transporter through an extended interface involving both TAP1 and TAP2 (Figure 3). CPXV012 remained positioned along the luminal/transmembrane-facing region of the transporter in all three simulations, although the precise orientation of its luminal segment differed between trajectories (Figure 3). This behavior is consistent with the RMSD, RMSF, and DSSP analyses, which showed that the luminal part of CPXV012 is more dynamic than the N-terminal/transmembrane helix (Figure 2; Figure S6; Figure S7).

**Figure 3.**
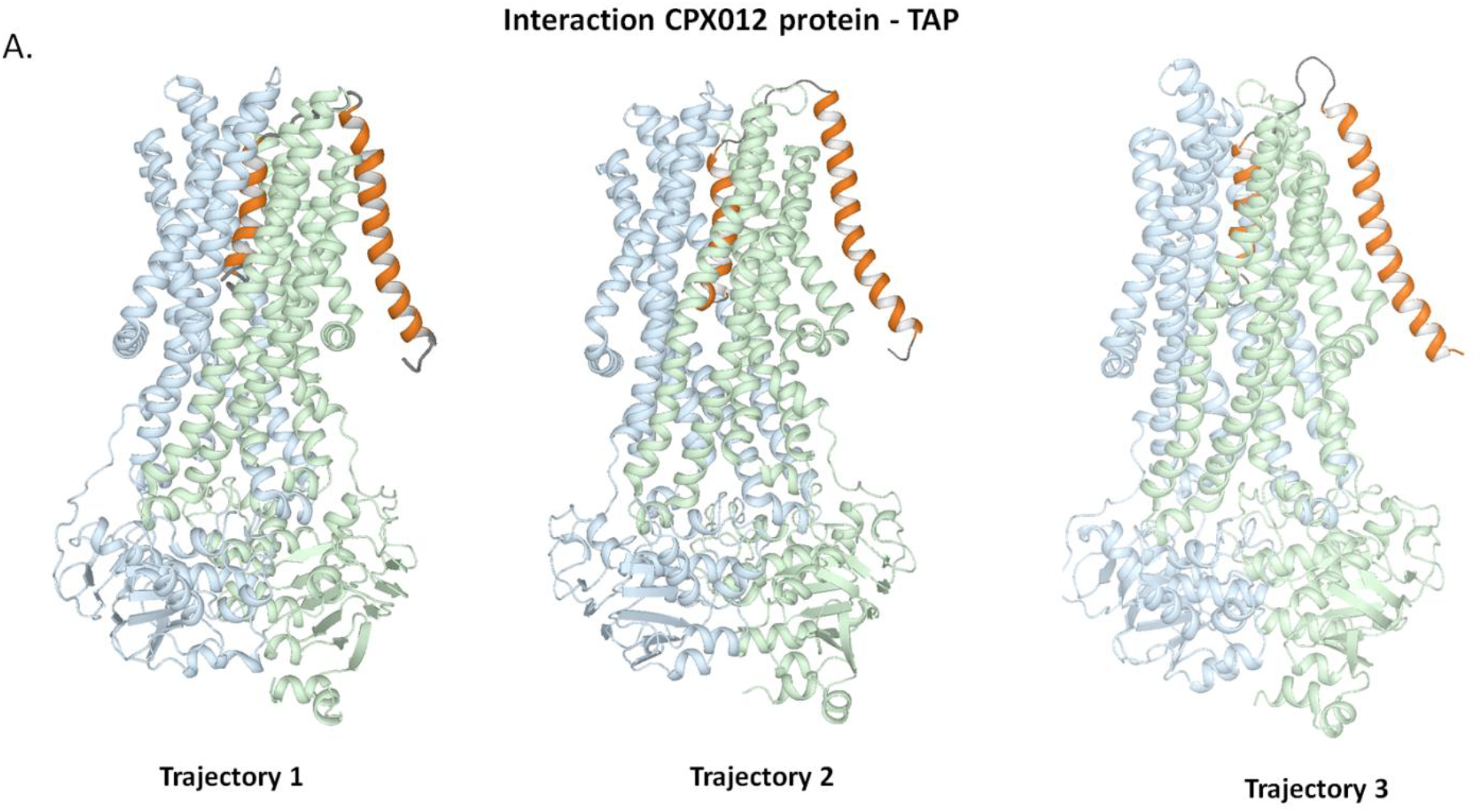
Model of the CPXV012–TAP complex in the open conformation. Representative final conformations of the CPXV012–TAP model obtained from three independent molecular dynamics trajectories. CPXV012 is shown in orange, TAP1 in green, and TAP2 in blue. The structures illustrate the position of CPXV012 relative to the open conformation of the TAP transporter.

The interaction network calculated from the combined trajectories showed that the CPXV012–TAP interface is dominated by recurrent contacts involving the ER-luminal region of CPXV012, particularly residues Y47– I69 (Figure S8). Both hydrogen bonds and salt bridges contributed to the interface (Figure S8). Several contacts involved charged CPXV012 residues, including R59, K64, and D66, which interacted with complementary charged or polar residues in TAP1 and TAP2. The presence of both TAP1- and TAP2-mediated interactions indicates that CPXV012 does not contact only one TAP subunit, but forms a distributed interface across the transporter (Figure 3; Figure S8).

The residue-pair interaction energy matrix identified several favorable CPXV012–TAP contacts (Figure S9). The strongest favorable terms included TAP2 R210–CPXV012 I69, TAP2 E417–CPXV012 K51, TAP1 D237–CPXV012 R59, TAP1 D288–CPXV012 K64, TAP1 R458–CPXV012 D66, TAP2 R273–CPXV012 I69, TAP1 E233–CPXV012 R59, TAP1 E292–CPXV012 K64, and TAP1 R185–CPXV012 E6 (Figure S9). These interactions were dominated by charge-complementary pairs and contacts involving the C-terminal I69-containing region of CPXV012.

Overall, the interface analysis suggests that the TAP-bound model of CPXV012 is stabilized by a combination of electrostatic and polar interactions, with the most prominent contacts involving the ER-luminal segment of CPXV012 (Figure S8; Figure S9). The concentration of favorable contacts in the Y47–I69 region is consistent with the proposed role of the CPXV012 luminal domain in TAP recognition and inhibition.

## Disscusion

This study provides a membrane-oriented structural model of CPXV012 and describes how its conformational properties may support interaction with TAP. The MD simulations indicate that CPXV012 maintains a two-helical organization in the open-conformation TAP model. The N-terminal/transmembrane helix remains comparatively stable across trajectories, whereas the ER-luminal region shows greater conformational variability. This distinction is important because previous functional studies assigned TAP inhibition primarily to the ER-luminal part of CPXV012 rather than to the transmembrane anchor itself (Luteijn et al., 2014; Lin et al., 2014). Luteijn et al. reported that CPXV012 is a type II membrane protein that blocks peptide transport by inhibiting ATP binding to TAP, and that the inhibitory activity is associated with its ER-luminal domain. Lin et al. further showed that CPXV012 acts after peptide binding and peptide-induced TAP conformational change, blocking ATP binding and hydrolysis through a lumenal trans-inhibition mechanism.

The present simulations are consistent with this functional framework. The stable N-terminal/transmembrane segment may serve mainly as a membrane-anchoring element that positions the viral protein relative to TAP. In contrast, the luminal region appears sufficiently structured to form a recurrent interface, but sufficiently flexible to adjust its contacts with TAP. This combination of local helicity and conformational adaptability may be important for a small viral membrane protein that must engage a large, dynamic ABC transporter. In this context, the increased mobility of the E37–K43 linker is particularly relevant, because this region connects the transmembrane anchor with the TAP-interacting luminal segment.

The interface analysis identifies a set of candidate interaction hotspots that may stabilize the CPXV012– TAP complex. In particular, the favorable contacts involving CPXV012 R59, K64, D66, I68, and I69 suggest that the C-terminal luminal region participates directly in TAP recognition. This observation agrees with the functional evidence that short C-terminal ER-luminal CPXV012 fragments can inhibit TAP activity. Lin et al. showed that the C-terminal 10 residues of CPXV012 are sufficient to inhibit peptide-stimulated ATP hydrolysis of TAP, with the peptide acting from the ER-luminal side in a trans-inhibition model. Therefore, the contacts observed here between the C-terminal Ile region of CPXV012 and TAP2 are mechanistically plausible, although their functional relevance requires direct mutational validation.

Recent cryo-EM data provide an important structural context for these findings. Lee et al. reported cryo-EM structures of TAP in complex with several viral inhibitors, including CPXV012, and concluded that structurally diverse viral inhibitors converge on a shared strategy of stalling TAP in its transport cycle (Lee et al., 2025). The RCSB entry for the TAP–CPXV012 complex describes TAP bound to CPXV012 in an outward-facing open state at 3.0 Å resolution and links this structure to the 2025 PNAS study by Lee, Manon, and Chen. The current MD analysis complements those static structural snapshots by showing that CPXV012 can preserve its global fold while allowing local rearrangements in the luminal inhibitory region.

The broader literature on viral TAP inhibition also supports an allosteric interpretation. Viral TAP inhibitors use different binding sites and structural strategies, but several ultimately prevent productive ATP-dependent peptide transport. Nolan et al. summarized that CPXV012 and HCMV US6 bind TAP from the ER-luminal side and block ATP binding, likely through distal allosteric effects, whereas other inhibitors such as EBV BNLF2a and HSV ICP47 interfere with TAP through different conformational states or substrate-binding steps. In this framework, the CPXV012–TAP contacts observed in the present model may represent structural elements that couple luminal binding of CPXV012 to impaired nucleotide-dependent TAP cycling.

The data also support the importance of the membrane environment for CPXV012 organization. The CD and NMR analyses showed that CPX-E1 and CPX-C2 are predominantly disordered in aqueous conditions but acquire ordered, mainly helix-like features in DPC micelles. This behavior is consistent with the idea that CPXV012 is not simply a soluble luminal peptide inhibitor, but a membrane-associated viral protein whose inhibitory region is spatially constrained by anchoring in or near the ER membrane. The NMR-derived structural restraints further support the presence of defined local structure in CPXV012-derived fragments under membrane-mimicking conditions.

Several limitations should be considered. First, the CPXV012–TAP interface described here is based on computational modeling and MD simulations, not on direct experimental mapping of contact residues. Second, DPC micelles provide a useful membrane-mimicking environment for NMR and CD studies, but they do not fully reproduce the physical properties of an ER lipid bilayer. Third, the peptide-fragment approach enables local structural analysis, but isolated fragments may not capture all long-range constraints present in full-length CPXV012. Finally, the strongest residue-pair interactions identified in the energy matrix should be treated as hypotheses for future testing rather than as confirmed functional determinants.

Future experiments should therefore focus on mutational validation of the predicted interface. Residues R59, K64, D66, I68, and I69 in CPXV012, together with the complementary TAP residues identified in the interaction matrix, are priority candidates for testing by peptide transport assays, ATP-binding/hydrolysis assays, and co-immunoprecipitation or crosslinking approaches. Simulations of additional TAP conformational states and ER-like lipid compositions would also help determine whether the observed interface is specific to the open-conformation model or persists across the TAP transport cycle.

In summary, the present results support a model in which CPXV012 combines a stable membrane-anchoring helix with a flexible but interaction-competent ER-luminal inhibitory region. This organization provides a structural rationale for how CPXV012 may engage TAP from the luminal side and stabilize an inhibited transporter state. The model is consistent with previous biochemical and structural studies of CPXV012-mediated TAP inhibition, while adding a dynamic view of the CPXV012–TAP interface.

## Acknowledgments

Computations were carried out using the computational resources of the Tricity Academic Supercomputer & Network Center of Informatics.

## Funding

This work was supported by the Polish National Science Centre grant no. UMO-2022/47/D/NZ7/02399 to N.K.

## Conflict of Interest

The authors declare that they have no competing financial interests or personal relationships that could have influenced the work reported in this paper.

## Data Availability

The data supporting the findings of this study are available from the corresponding author upon reasonable request.

## CRediT author statement

Natalia Karska: Conceptualization, Methodology, Investigation, Formal analysis, NMR structure determination, Writing – original draft, Writing – review and editing, Supervision, Project administration, Funding acquisition. Behnaz Mizraeli: Formal analysis, NMR data analysis, NMR structure determination, Magdalena J. Ślusarz: Methodology, Software, Formal analysis, Investigation, Visualization, full-length CPXV012 modeling, molecular modeling of the CPXV012–TAP system, Writing – review and editing. Przemysław Karpowicz: Investigation, Resources, peptide synthesis, peptide purification, Igor Zhukov: Methodology, Investigation, NMR data acquisition, NMR spectroscopy, structure generation, Data curation, Writing – review and editing. Sylwia Rodziewicz-Motowidło: Supervision, Resources, Writing – review and editing.

## Materials and Methods

### Chemical peptide synthesis and purification

Peptides related to the (an ER-luminal domain fragment of CPXV012 (E.CPXV), a transmembrane fragment of CPXV012(T.CPXV012) and cytoplasmic fragment of CPXV012 (C. CPXV012)) were synthesized by solid-phase synthesis technique (Millipore 9050 Plus PepSynthesizer, Millipore Corporation, Burlington, VT, USA)) using the Fmoc chemistry strategy at Rink Amide resin (capacity 0.2 mmol/g, Rapp Polymere GmbH. Germany) [20,21]. In addition to the individual peptide fragments, the full-length CPXV protein was synthesized using the same approach. Fmoc-protected amino acids and coupling reagents were purchased from NovaBiochem (Nottingham, UK) or Sigma-Aldrich (Poznań, Poland). To avoid unwanted oxidation of the sulfhydryl group of cysteine and disulfide bond formation of the methionine during synthesis and purification, cysteine and methionine residues were replaced with their isosteric analogues, (S)-2-aminobutyric acid (Abu) and norleucine (Nle), respectively. The peptides were cleaved from the resin using a cleavage mixture composed of trifluoroacetic acid, triisopropylsilane, phenol, and water (95:2.5:2.5) at room temperature for 4 h with gentle vibration. The exhausted resin was removed by filtration, and the filtrate was concentrated under reduced pressure. Crude peptides were precipitated by trituration with cold diethyl ether, collected by centrifugation at 4000 rpm for 10 min, and washed repeatedly to remove residual scavengers (the process was repeated three times).

Purification of peptide fragments was performed by reversed-phase high-performance liquid chromatography (RP-HPLC) using a system equipped with two K1001 pumps (Knauer, Germany) and a UV–Vis detector coupled to a Gilson fraction collector. Depending on peptide hydrophobicity, separations were carried out on Kromasil C8 or C4 columns (10 × 250 mm, particle size 5 μm, pore size 300 Å). Linear gradients of acetonitrile– water or methanol–water mixtures containing either 0.1% (v/v) trifluoroacetic acid or 0.1 M triethylammonium phosphate (TEAP, pH 3.0) were used as eluents at a flow rate of 5 mL/min. Peptide elution was monitored at 223 nm. The purity of all synthesized peptides was higher than 98%, as determined by analytical RP-HPLC. Molecular masses were confirmed by matrix-assisted laser desorption/ionization time-of-flight mass spectrometry (MALDI-TOF MS) using a Bruker Biflex III spectrometer (Bruker Daltonics, Germany) with α-cyano-4-hydroxycinnamic acid as the matrix, as well as by electrospray ionization mass spectrometry (ESI-MS) recorded on a Shimadzu IT-TOF mass spectrometer (Shimadzu, Japan).

### CD spectropolarimetry

CD studies were performed in DPC micelles (5 mM and/or 100 mM, POCH, Poland) in water (pH 7.0) or PBS (pH 7.4). The peptide solutions in the DPC micelles were prepared according to standard procedures. The CD spectra were recorded for each peptide (0.15 mg/mL) in the range of 185–260 nm with a Jasco J-815 spectropolarimeter (Jasco, Easton, USA). For each peptide, the CD spectra were recorded three times, at 30 °C using a 1-mm cell, and are shown as the mean residue molar ellipticity (MRME, degree×cm^2^×dmol^™1^) versus wavelength λ (nm). CD spectra were recorded for peptide solutions at a concentration of 0.15 mg/mL in the wavelength range of 185–260 nm at 30 °C using a quartz cuvette with a 1 mm path length using a Jasco J-815 spectropolarimeter (Jasco, Easton, MA, USA). Measurements were performed For each peptide, three independent spectra were collected and averaged. The data were expressed as mean residue molar ellipticity (MRME, deg·cm^2^·dmol^−1^) as a function of wavelength.

### NMR Spectroscopy

The NMR samples were prepared by dissolving 1.5 mg peptides in 100 mM DPC-d38 micelle solutes in H2O/D2O 90%/10%. All NMR experiments were performed at 308 K on Varian Inova 500 (1H resonance frequency 500.606 MHz). Spectrometer equipped with three channels, z-gradient unit, and 1H/13C/15N triple resonance probehead with inverse detection. The collected NMR data contained the homonuclear 2D NMR experiments – 1H-1H TOCSY data recorded with mixing time 80 ms, and 1H-1H NOESY acquired with mixing time 150 ms. The heteronuclear 2D NMR experiments include 1H-13C HSQC and 1H-15N HSQC detected on natural abundance of 13C and 15N isotopes. All chemical shifts were referenced with respect to external sodium 2,2-dimethyl-2-silapentane-5-sulfonate (DSS) using Ξ = 0.251449530 and 0.101329118 ratios for indirectly referenced 13C and 15N resonances, re-spectively (Wishart et al., 1995). Recorded NMR data were processed by NMRPipe software (Delaglio et al., 1995) and analyzed with the NMRFAM-Sparky program (Lee et al., 2015).

### Evaluation of the 3D structure

Assignments of the 1H, 13C, and 15N resonances were achieved with jointedanalysis 2D homonuclear (TOCSY, NOESY) and heteronuclear (1H-13C, and 1H-15N HSQC) data sets. Initial 3D structures of CPX-E and CPX-C2 peptides were established with CYANA (version 3.98.15) software (Güntert, 2004). There are 178 for CPX-E1 (83 intraresidue, 51 sequential, 12 medium, and 32 long-range) and 103 for CPX-C2 (51 intraresidue, 29 sequential, and 23 medium-range) distance constraints were yielded by inspection of NOESY spectrum acquired with mixing time 150 ms. These data were supplemented with 44 (CPX-E1) or 20 (CPX-C2) restraints for the backbone ϕ, ψ torsion angles deduced from analysis assigned 1H, 13C, and 15N resonances with TALOSn software (Shen and Bax, 2013). In addition, the 20 (CPX-E1) or 8 (CPX-C2) constraints for hydrogen bonds were established basing on geo-metric criteria and applied, as rHN −O = 2.1 ± 0.5 Å, and rN −O = 3.0 ± 0.5 Å, at refine stage of calculations. Topology of the NLE and ABU residues were initially created with MolMol software (Koradi et al., 1996) and transformed to cyana library. The final refinement in explicit solution were performed with Yasara (version 25.1.13) software (Krieger et al., 2002) utilizing modified versions of nmr_refine.mcr, and nmr_setdefault.mcr macros included in Yasara library (Krieger and Vriend, 2015).

## Notes

### Competing Interest Statement

The authors have declared no competing interest.

