## Supplementary material for "Membrane-dependent structural organization of cowpox virus CPXV012 and its recognition of TAP": Table_Suplementary

**Table S1. Structural restraints and quality assessment of the NMR-derived CPX-E1 and CPX-C2 models.** The table summarizes the number and distribution of NOESY-derived distance restraints, hydrogen-bond restraints, torsion angle restraints, RMSD values, Ramachandran statistics, and RMS Z-scores used to evaluate the quality of the calculated structural ensembles for CPX-E1 and CPX-C2.

| | ***CPX-E1*** | ***CPX-C2*** |
|---|---|---|
| *NOESY distance constraints* | | |
| Intraresidue | *83* | *51* |
| Sequential *($\|i-j\| = 1$)* | *51* | *29* |
| Medium range *($\|i-j\| \leq 5$)* | *12* | *23* |
| Long range *($\|i-j\| > 5$)* | *32* | *0* |
| Hydrogen bonds | *10* | *4* |
| *Torsion angle restraints*[1] | *44* | *20* |
| Backbone angles *(φ/ψ)* | *9* | *5* |
| *RMSD to main structure* | | |
| RMSD backbone (Å) | 1.15 ± 0.30 | 0.29 ± 0.12 |
| RMSD heavy atoms (Å) | 1.23 ± 0.37 | 1.27 ± 0.24 |
| *Ramachandran plot*[2] | | |
| Residues in most favored regions (%) | 84.1 | 100.0 |
| Residues in additionally allowed regions (%) | z15.4 | *0* |
| Residues in generously allowed regions (%) | *0.4* | *0* |
| *RMS Z-score*[3] | | |
| Bond lengths *(Å)* | -0.123 ± 0.094 | -0.474 ± 0.103 |
| Bond angles | -1.229 ± 0.110 | 0.777 ± 0.128 |
| Dihedral angles | 1.336 ± 0.180 | 2.331 ± 0.424 |
| Side chains planarity | 0.641 ± 0.042 | 0.642 ± 0.073 |
| Nonbonded interactions | -2.035 ± 0.520 | -2.293 ± 0.527 |
| Model quality | -3.406 ± 0.363 | -2.870 ± 0.388 |
